# Transcriptional landscape of DNA repair genes underpins a pan-cancer prognostic signature associated with cell cycle dysregulation and tumor hypoxia

**DOI:** 10.1101/519603

**Authors:** Wai Hoong Chang, Alvina G. Lai

## Abstract

Overactive DNA repair contributes to therapeutic resistance in cancer. However, pan-cancer comparative studies investigating the contribution of *all* DNA repair genes in cancer progression employing an integrated approach have remained limited. We performed a multicohort retrospective analysis to determine the prognostic significance of 138 DNA repair genes in 16 cancer types (n=16,225). Cox proportional hazards analyses revealed a significant variation in the number of prognostic genes between cancers; 81 genes were prognostic in clear cell renal cell carcinoma while only two genes were prognostic in glioblastoma. We reasoned that genes that were commonly prognostic in highly correlated cancers revealed by Spearman’s correlation analysis could be harnessed as a molecular signature for risk assessment. A 10-gene signature, uniting prognostic genes that were common in highly correlated cancers, was significantly associated with overall survival in patients with clear cell renal cell (P<0.0001), papillary renal cell (P=0.0007), liver (P=0.002), lung (P=0.028), pancreas (P=0.00013) or endometrial (P=0.00063) cancers. Receiver operating characteristic analyses revealed that a combined model of the 10-gene signature and tumor staging outperformed either classifiers when considered alone. Multivariate Cox regression models incorporating additional clinicopathological features revealed that the signature was an independent predictor of overall survival. Tumor hypoxia is associated with adverse outcomes. Consistent across all six cancers, patients with high 10-gene and high hypoxia scores had significantly higher mortality rates compared to those with low 10-gene and low hypoxia scores. Functional enrichment analyses revealed that high mortality rates in patients with high 10-gene scores were attributable to an overproliferation phenotype. Death risk in these patients was further exacerbated by concurrent mutations of a cell cycle checkpoint protein, *TP53*. The 10-gene signature identified tumors with heightened DNA repair ability. This information has the potential to radically change prognosis through the use of adjuvant DNA repair inhibitors with chemotherapeutic drugs.

## Introduction

Genetic material must be transmitted in its original, unaltered form during cell division. However, DNA faces continuous assaults from both endogenous and environmental agents contributing to the formation of permanent lesions and cell death. To overcome DNA damage threats, living systems have evolved highly coordinated cellular machineries to detect and repair damages as they occur. However, DNA repair mechanisms and consequently DNA damage responses (DDR) are often deregulated in cancer cells and such aberrations may contribute to cancer progression and influence prognosis. Overexpression of DNA repair genes allow tumor cells to overcome the cytotoxic effects of radiotherapy and chemotherapy. As such, inhibitors of DNA repair can increase the vulnerability of tumor cells to chemotherapeutic drugs by preventing the repair of deleterious lesions^1^.

There are six main DNA repair pathways in mammalian cells. Single-strand DNA damage are repaired by the base excision repair (BER), nucleotide excision repair (NER) and mismatch repair (MR) pathways. The poly(ADP-ribose) polymerase (PARP) gene family encodes key players of the BER pathway involved in repairing damages induced by ionizing radiation and alkylating agents^2,3^. Replication errors are corrected by the MR pathway while the NER pathway is responsible for removing bulky intercalating agents^4,5^. Tumor cells with deficiencies in the NER pathway have increased sensitivity to platinum-based chemotherapeutic drugs (cisplatin, oxaliplatin etc.)^6,7^. Double-strand breaks induced by ionizing radiation are more difficult to repair and thus are highly cytotoxic. Dysregulation of genes involved in the homology-directed repair (HDR), non-homologous end joining (NHEJ) and Fanconi anemia (FA) pathways are associated with altered repair of double-strand breaks.

Aberrations in DNA repair genes are widespread in most cancers; hence they represent attractive candidates for pharmacological targeting to improve radiosensitivity and chemosensitivity^8^. In a process known as ‘synthetic lethality’, faults in two or more DNA repair genes or pathways together would promote cell death, while defects in a single pathway may be tolerated^1^. Functional redundancies in repair pathways allow tumor cells to rely on a second pathway for repair in the event that the first pathway is defective. Based on the principles of synthetic lethality, inhibition of the second pathway will confer hypersensitivity to cytotoxic drugs in cells with another malfunctioning pathway. This promotes cell death because DNA lesions can no longer be repaired by either pathway. For instance, PARP inhibitors (targeting the BER pathway) could selectively kill tumor cells that have *BRCA1* or *BRCA2* mutations (defective HDR pathway) while not having any toxic effects on normal cells^9,10^.

Since one DDR pathway could compensate for another, there is a need for a pan-cancer, large-scale, systematic study on *all* DNA repair genes to reveal similarities and differences in DDR signaling between cancer types, which is limited at present. In this study, we explored pan-genomic expression patterns of 138 DNA repair genes in 16 cancer types. We developed and validated the prognostic significance of a 10-gene signature that can be used for rapid risk assessment and patient stratification. There are considerable variations in the success of chemotherapy and radiotherapy regimes between cancer types. Such differences may be explained by the complex cancer-specific nature of DDR defects. Prognostic biomarkers of DNA repair genes are needed to allow the use of repair inhibitors in a stratified, non-universal approach to expose the selective vulnerabilities of tumors to therapeutic agents.

## Materials and methods

A list of 138 DNA repair genes is available in Table S1.

### Study cohorts

We obtained RNA-sequencing datasets for the 16 cancers from The Cancer Genome Atlas (TCGA)^11^ (n=16,225) (Table S2). TCGA Illumina HiSeq rnaseqv2 Level 3 RSEM normalized data were retrieved from the Broad Institute GDAC Firehose website. Gene expression profiles for each cancer types were separated into tumor and non-tumor categories based on TCGA barcodes and converted to log_2_(x + 1) scale. To compare the gene-by-gene expression distribution in tumor and non-tumor samples, violin plots were generated using R. The nonparametric Mann-Whitney-Wilcoxon test was used for statistical analysis.

### Calculation of 10-gene scores and hypoxia scores

The 10-gene scores for each patient were determined from the mean log_2_ expression values of 10 genes: *PRKDC, NEIL3, FANCD2, BRCA2, EXO1, XRCC2, RFC4, USP1, UBE2T* and *FAAP24).* Hypoxia scores were calculated from the mean log_2_ expression values of 52 hypoxia signature genes^12^. For analyses in Figure 5, patients were delineated into four categories using median 10-gene scores and hypoxia scores as thresholds. The nonparametric Spearman’s rank-order correlation test was used to determine the relationship between 10-gene scores and hypoxia scores.

### Differential expression analyses comparing expression profiles of high-score and low-score patients

Patients were median dichotomized into low- and high-score groups based on their 10-gene scores in each cancer type. Differential expression analyses were performed using the linear model and Bayes method executed by the limma package in R. P values were adjusted using the Benjamini-Hochberg false discovery rate procedure. We considered genes with log_2_ fold change of > 1 or < −1 and adjusted P-values < 0.05 as significantly differentially expressed between the two patient groups.

### Functional enrichment and pathway analyses

To determine which biological pathways were significantly enriched, differentially expressed genes were mapped against the Gene Ontology (GO) and Kyoto Encyclopedia of Genes and Genomes (KEGG) databases using GeneCodis^13^. The Enrichr tool was used to investigate transcription factor protein-protein interactions that were associated with the differentially expressed genes^14,15^.

### Survival analysis

Univariate Cox proportional hazards regression analyses were performed using the R survival and survminer packages to determine if expression levels of individual DNA repair genes as well as those of the 10-gene scores were significantly associated with overall survival. Multivariate Cox regression was employed to determine the influence of additional clinical variables on the 10-gene signature. Hazard ratios (HR) and confidence intervals were determined from the Cox models. HR greater than one indicated that a covariate was positively associated with even probability or increased hazard and negatively associated with survival duration. Non-significant relationship between scaled Schoenfeld residuals supported the proportional hazards assumption in the Cox model. Both survival and survminer packages were also used for Kaplan-Meier analyses and log-rank tests. For Kaplan-Meier analyses, patients were median dichotomized into high- and low-score groups using the 10-gene signature. To determine the predictive performance (specificity and sensitivity) of the signature in relation to tumor staging parameters, we employed the receiver operating characteristic (ROC) analysis implemented by the R survcomp package, which also calculates area under the curve (AUC) values. AUC values can fall between 1 (perfect marker) and 0.5 (uninformative marker).

### *TP53* mutation analysis

TCGA mutation datasets (Level 3) were retrieved from GDAC Firehose to annotate patients with mutant *TP53.* To ascertain the association of *TP53* mutation with the 10-gene signature on overall survival, we employed the Kaplan-Meier analysis and log-rank tests implemented in R.

All plots were generated using R pheatmap and ggplot2 packages^16^. Venn diagram was generated using the InteractiVenn tool^17^.

## Results

### Prognosis of DNA repair genes in 16 cancer types and the development of a 10-gene signature

A total of 187 genes associated with six DDR pathways found in mammalian cells were curated: BER (33 genes), MR (23 genes), NER (39 genes), HDR (26 genes), NHEJ (13 genes) and FA (53 genes)^18^ (Fig. 1, Table S1). Of the 187 genes, 49 were represented in two or more pathways, yielding 138 non-redundant candidates. To determine which of the 138 DNA repair genes conferred prognostic information, we employed Cox proportional hazards regression on all genes individually on 16 cancer types to collectively include 16,225 patients^11^ (Table S2). In clear cell renal cell carcinoma, 81 genes were found to be significantly associated with overall survival; this cancer had the highest number of prognostic DNA repair genes (Table S3). This is followed by 54, 53, 46, 44 and 33 prognostic genes in cancers of the pancreas, papillary renal cell, liver, lung and endometrium respectively (Table S3). In contrast, cancers of the brain (glioblastoma: 2 genes), breast (5 genes), cervix (6 genes) and esophagus (7 genes) had some of the lowest number of prognostic DNA repair genes (Table S3), suggesting that there is a significant degree of variation in the contribution of DNA repair genes in predicting survival outcomes. Spearman’s rank-order correlation analysis revealed a hub of five highly correlated cancers (lung, papillary renal cell, pancreas, liver and endometrium), indicating that a good number of prognostic DNA repair genes were shared between these cancers (Spearman’s rho=0.21 to 0.44) (Fig. S1). We rationalized that prognostic genes that are common in these highly correlated cancers could form a new multigenic risk assessment classifier. Ten genes were prognostic in the five highly correlated cancers: *PRKDC* (NHEJ), *NEIL3* (BER), *FANCD2* (FA), *BRCA2* (HDR and FA), *EXO1* (MR), *XRCC2* (HDR), *RFC4* (MR and NER), *USP1* (FA), *UBE2T*(FA) and *FAAP24* (FA), which, interestingly, represent members from all six DDR pathways.

**Figure 1.**
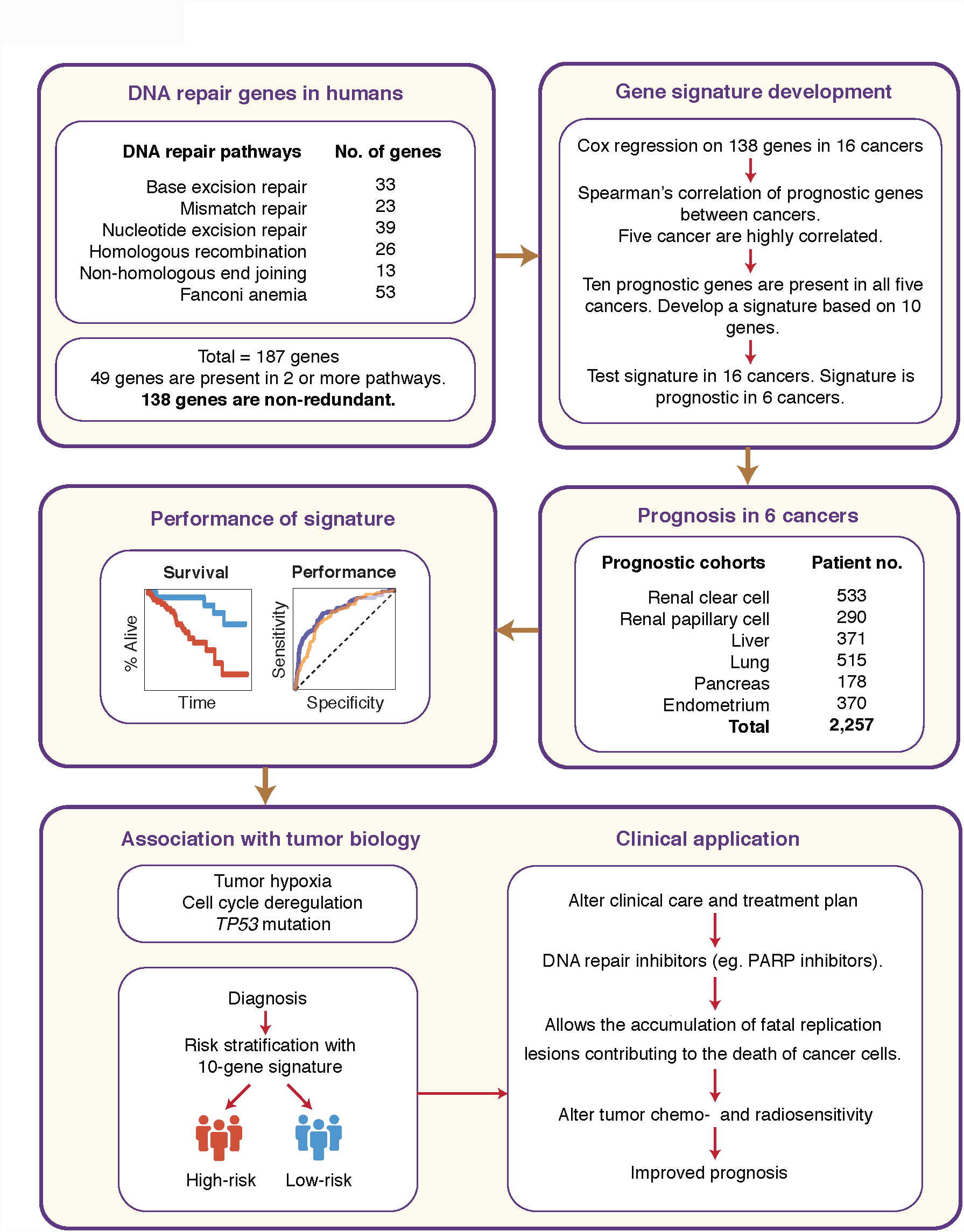
Schematic representation of the study design and development of the 10-gene signature. DNA repair genes from six major pathways were manually curated to generate a non-redundant list containing 138 genes. Cox proportional hazards regression was employed to determine the significance of each individual genes in predicting overall survival in 16 cancer types. Spearman’s correlation analyses revealed that five cancer types exhibited a high degree of correlation in terms of their prognostic genes. Ten genes were found to be prognostic in all five cancers; these genes subsequently formed the 10-gene signature. The ability of the signature in predicting survival outcomes was tested using Kaplan-Meier, Cox regression and receiver operating characteristic methods. The signature could predict high-risk patients in six cancer types (n=2,257). Associations of the signature with tumor hypoxia, cell cycle deregulation and *TP53* mutation were investigated. Potential clinical applications of the signature were proposed.

### A 10-gene signature predictive of DDR signaling is an independent prognostic classifier in 6 cancer types

The aforementioned ten genes were employed as a new prognostic model to evaluate whether they were significantly associated with overall survival in all 16 cancer types. A 10-gene score for each patient was calculated by taking the mean expression of all ten genes. Patients were median dichotomized based on their 10-gene scores into a low- and high-score groups. The 10-gene signature could predict patients at significantly higher risk of death in the five cancers that were originally highly correlated (Fig. S1), and in one additional cancer (clear cell renal cell carcinoma) (Fig. 2). Kaplan-Meier analyses demonstrated that patients categorized within high-score groups had significantly poorer survival rates: clear cell renal cell (log-rank P<0.0001), papillary renal cell (P=0.0007), liver (P=0.002), lung (P=0.028), pancreas (P=0.00013) and endometrium (P=0.00063) (Fig. 2). Expression profiles of the 10 genes in tumor and non-tumor samples showed a general distribution that were comparable among the six cancer types. Mann-Whitney-Wilcoxon tests revealed that a vast majority of genes were significantly upregulated in tumor samples with a few minor exceptions (Fig. S2). *USP1* was significantly downregulated in tumors of papillary renal cell and endometrium (Fig. S2). Only four non-tumor samples were available in the pancreatic cancer cohort, precluding robust statistical analyses. Due to limitations in sample size, only *UBE2T* was observed to be significantly upregulated in pancreatic tumors (Fig. S2).

**Figure 2.**
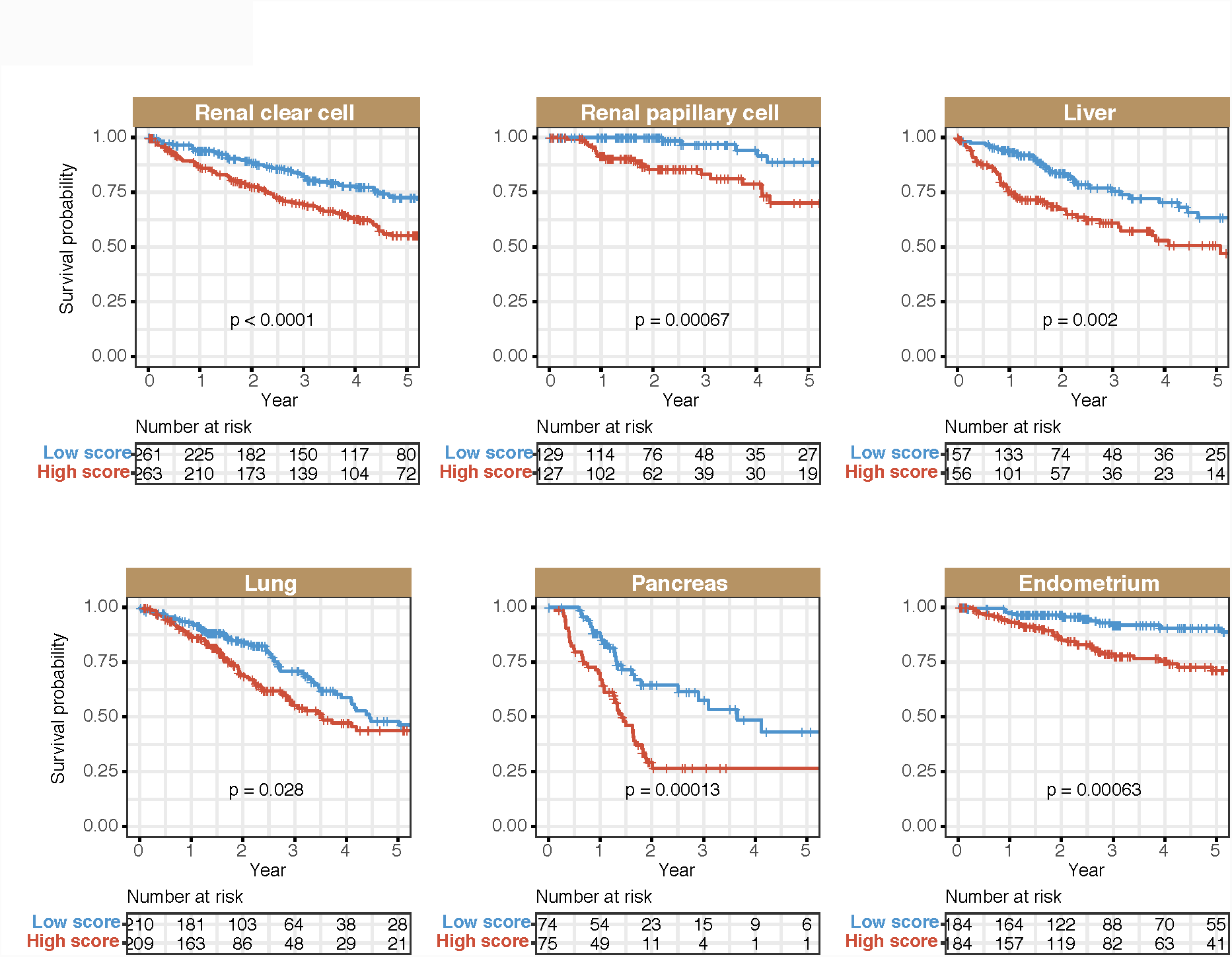
Patient stratification using the 10-gene signature in six cancer types. Kaplan-Meier analyses of overall survival on patients stratified into high- and low-score groups using the 10-gene signature. P values were determined from the log-rank test.

To evaluate the independent predictive value of the signature over the current tumor, node and metastasis (TNM) staging system, we applied the signature on patients separated by TNM stage: early (stages 1 and/or 2), intermediate (stages 2 and/or 3) and late (stages 3 and/or 4) disease stages. Remarkably, the signature successfully identified high risk patients in early (liver, lung, pancreas, endometrium), intermediate (papillary renal cell, liver, pancreas, endometrium) and late (clear cell renal cell, papillary renal cell, liver, endometrium) TNM stages (Fig. 3). Collectively, this implied that the signature offered an additional resolution of prognosis within similarly staged tumors and that the signature retained excellent prognostic ability in individual tumor groups when considered separately.

**Figure 3.**
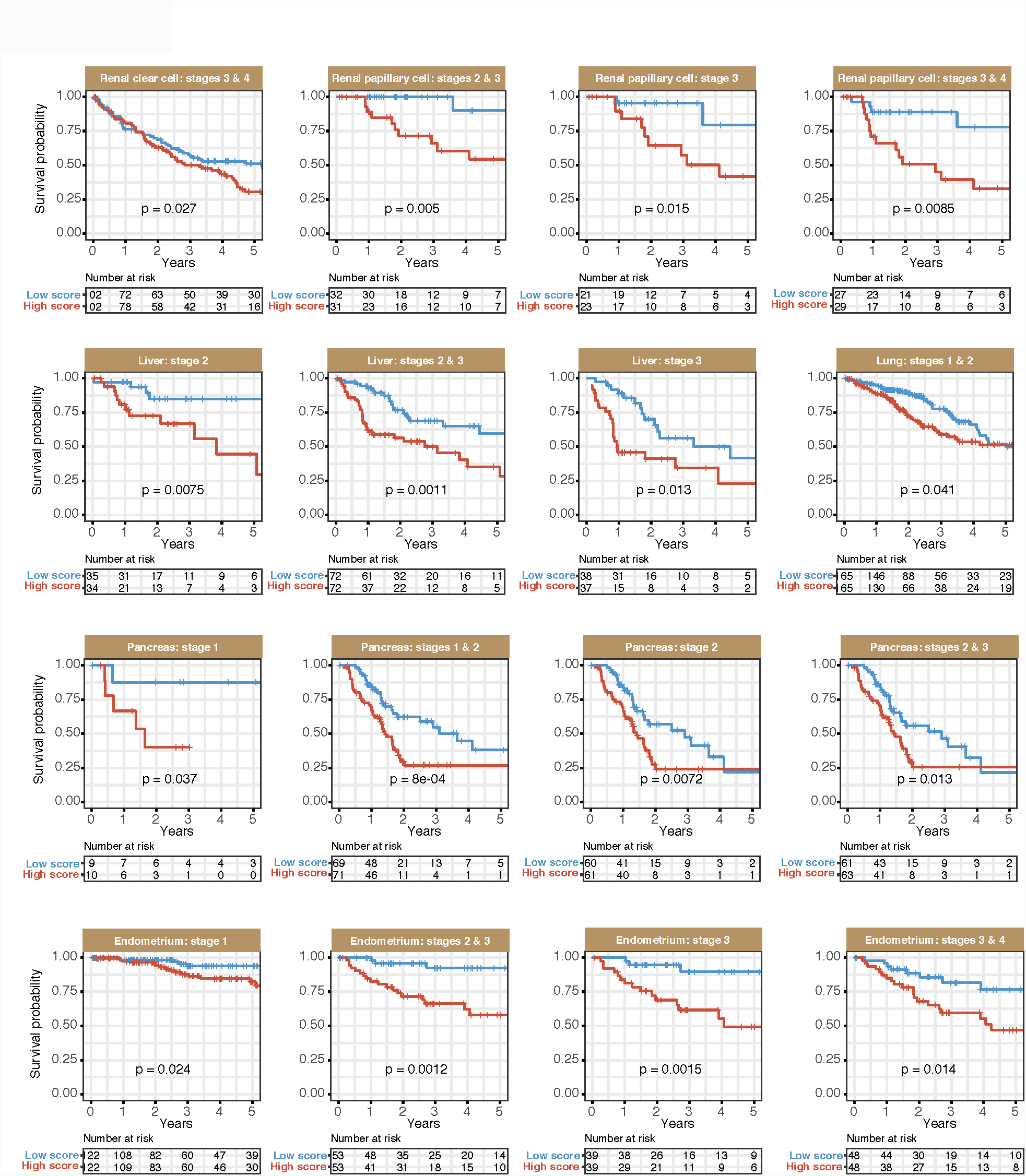
Independence of the 10-gene signature over TNM staging. Kaplan-Meier analyses were performed on patients categorized according to tumor TNM stages that were further stratified using the 10-gene signature. The signature successfully identified patients at higher risk of death in all TNM stages. P values were determined from the log-rank test. TNM: tumor, node, metastasis.

To evaluate the predictive performance of the 10-gene signature on 5-year overall survival, we employed receiver operating characteristic (ROC) analyses on all six cancers. Comparing the sensitivity and specificity of the signature in relation to TNM staging revealed that the signature outperformed TNM staging in cancers of the papillary renal cell (AUC=0.832 vs. AUC=0.640), pancreas (AUC=0.697 vs. AUC=0.593) and endometrium (AUC=0.700 vs. AUC=0.674) (Fig. 4). Importantly, when the signature was used in conjunction with TNM staging as a combined model, its performance was superior to either classifiers when they were considered individually: clear cell renal cell (AUC=0.792), papillary renal cell (AUC=0.868), liver (AUC=0.751), lung (AUC=0.693), pancreas (AUC=0.698) and endometrium (AUC=0.764) (Fig. 4).

**Figure 4.**
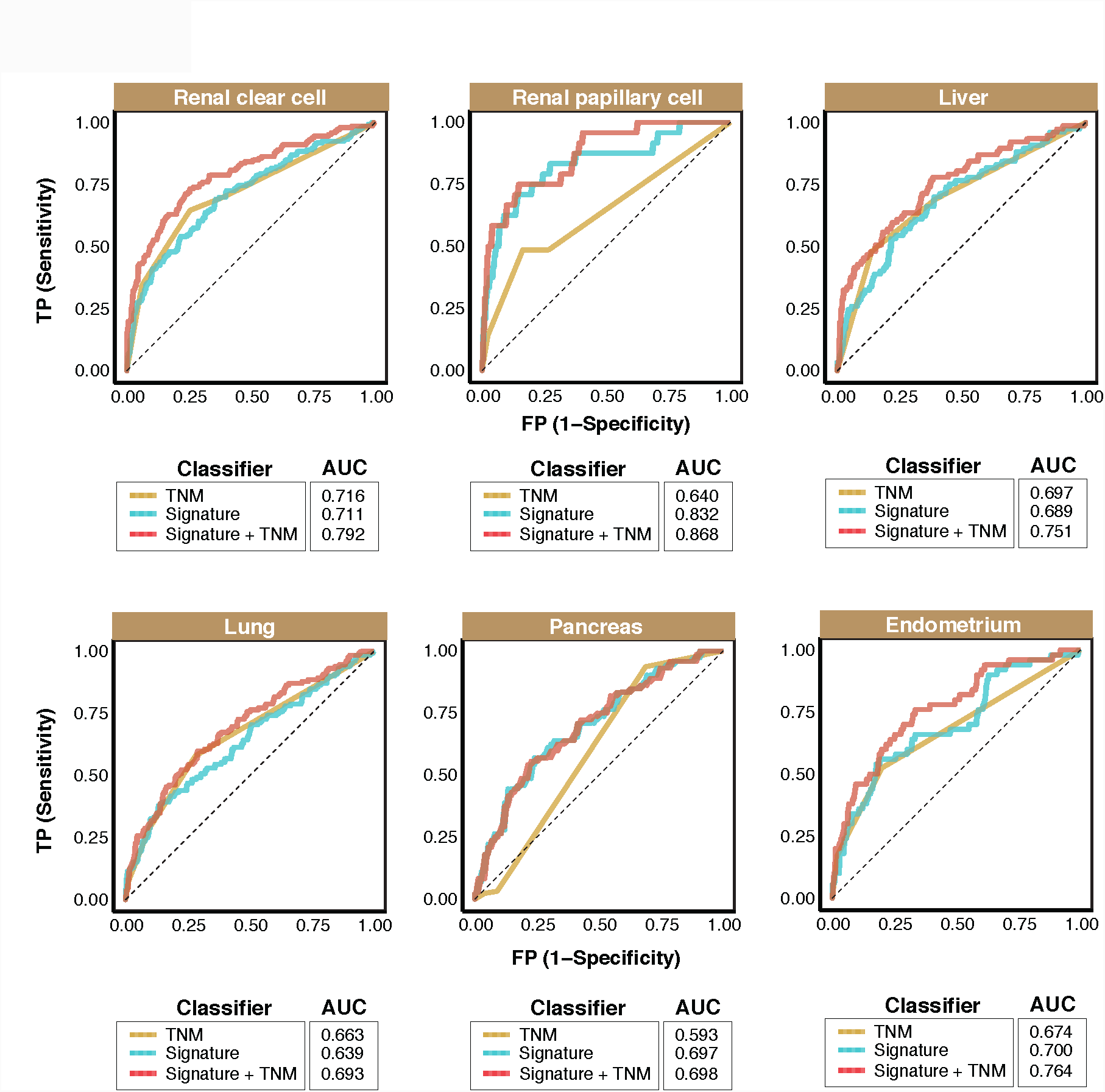
Predictive performance of the 10-gene signature. Receiver operating characteristic (ROC) was employed to determine the specificity and sensitivity of the signature in predicting 5-year overall survival in all six cancer types. ROC curves generated based on the 10-gene signature, TNM staging and a combination of 10-gene signature and TNM staging were depicted. AUC: area under the curve. TNM: tumor, node, metastasis. AUCs for TNM staging were in accordance with previous publications employing TCGA datasets^19,20^.

We next employed multivariate Cox regression models to examine whether the association between high 10-gene scores and increased mortality was not due to underlying clinical characteristics of the tumors. Univariate analysis revealed that TNM staging is not prognostic in pancreatic cancer (hazard ratio [HR]=1.339, P=0.153), hence this cancer was excluded from the multivariate model involving TNM (Table 1). For the five remaining cancer types, even when TNM staging was considered, the signature significantly distinguished survival outcomes in high-versus low-score patients, confirming that it is an independent prognostic classifier: clear cell renal cell (HR=1.555, P=0.0058), papillary renal cell (HR=1.677, P=0.032), liver (HR=1.650, P=0.029), lung (HR=1.301, P=0.032) and endometrium (HR=2.113, P=0.013) (Table 1).

### Crosstalk between DDR signaling and tumor hypoxia

Tumor hypoxia is a well-known barrier to curative treatment. It is often associated with poor prognosis^19,20^, which may be a result of tumor resistance to chemotherapy and radiotherapy^21,22^. Since both the upregulation of DNA repair genes and hypoxia are linked to therapeutic resistance, we rationalized that incorporating hypoxia information in the 10-gene signature would allow further delineation of patient risk groups. Patients with high 10-gene scores had significantly poorer survival outcomes and we predict that these patients have tumors that are more hypoxic, and that oxygen deprivation could influence DDR signaling to enhance tumor resistance to apoptotic stimuli leading to more aggressive disease states. We calculated hypoxia scores for each patient using a mathematically derived hypoxia gene signature consisting of 52 genes^12^. Hypoxia scores were defined as the mean expression of the 52 genes. Patients for each of the six cancer types were divided into four categories using the median 10-gene and hypoxia scores: 1) high scores for both 10-gene and hypoxia, 2) high 10-gene and low hypoxia scores, 3) low 10-gene and high hypoxia scores and 4) low scores for both 10-gene and hypoxia (Fig. 5A). Remarkably, significant positive correlations were observed between 10-gene scores and hypoxia scores consistent across all six cancer types: clear cell renal cell (rho=0.363, P<0.0001), papillary renal cell (rho=0.518, P<0.0001), liver (rho=0.615, P<0.0001), lung (rho=0.753, P<0.0001), pancreas (rho=0.582, P<0.0001) and endometrium (rho=0.527, P<0.0001) (Fig. 5A). This suggests that tumor hypoxia may influence DDR signaling and potentially, patient outcomes.

We generated Kaplan-Meier curves and employed the log-rank test to determine whether there were differences in overall survival outcomes among the four patient groups. Combined relation of hypoxia and 10-gene scores revealed significant associations with overall survival in all six cancers (Fig. 5B). Patients classified within the ‘high 10-gene and high hypoxia’ category had significantly poorer survival rates compared to those with low 10-gene and low hypoxia scores: clear cell renal cell (HR=2.316, P<0.0001), papillary renal cell (HR=7.635, P=0.0011), liver (HR=2.615, P=0.00013), lung (HR=1.832, P=0.0021), pancreas (HR=2.680, P=0.00079) and endometrium (HR=2.707, P=0.0075) (Table 2; Fig. 5B). Our results suggest that the combined effects of hypoxia and heightened expression of DNA damage repair genes may be linked to tumor progression and increased mortality risks. It remains unknown in this context whether the basis for differential sensitivity to chemotherapy would be explained, in part, by DNA repair ability of tumor cells exposed to chronic hypoxia environments.

**Figure 5.**
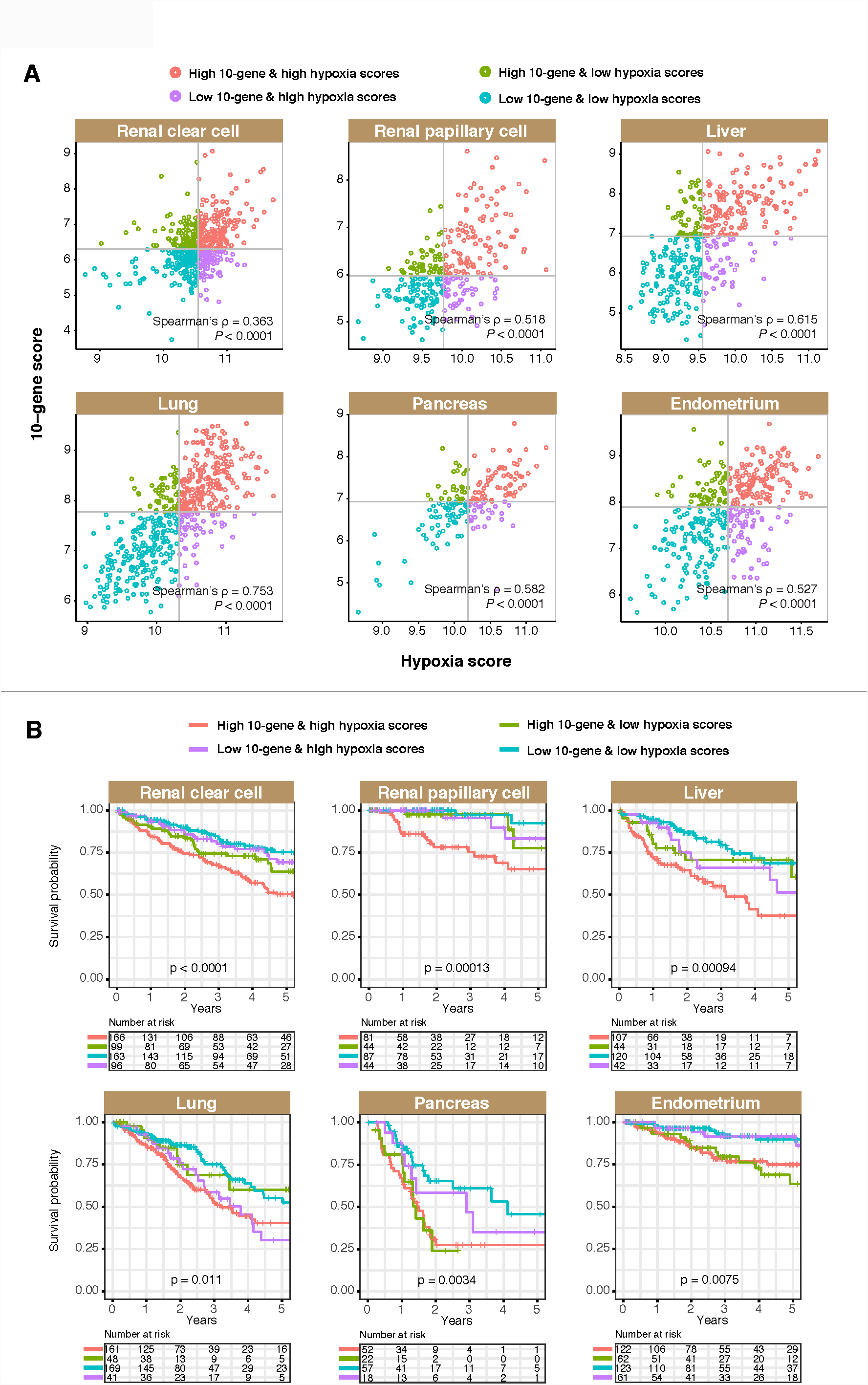
Association between the 10-gene signature and tumor hypoxia. **(A)** Scatter plots depict significant positive correlation between 10-gene scores and hypoxia scores in all six cancers. Patients were color-coded and separated into four categories based on their 10-gene and hypoxia scores. **(B)** Kaplan-Meier analyses were performed on the four patient categories to assess the effects of combined relationship of hypoxia and the signature on overall survival.

### Patients with high 10-gene scores had an overproliferation phenotype due to cell cycle dysregulation

The cell cycle represents a cellular gatekeeper that controls how cells grow and proliferate. Cyclins and cyclin-dependent kinases (CDKs) allow cells to progress from one cell cycle stage to the next; a process that is antagonized by CDK inhibitors. Many tumors overexpress cyclins or inactivate CDK inhibitors, hence resulting in uncontrolled cell cycle entry, loss of checkpoint and uninhibited proliferation^23-25^. Targeting proteins responsible for cell cycle progression would thus be an attractive measure to limit tumor cell proliferation. This has led to the development of numerous CDK inhibitors as anticancer agents^26,27^. DNA repair is tightly coordinated with cell cycle progression. Certain DNA repair mechanisms are dampened in nonproliferating cells, while repair pathways are often perturbed during tumor development. Perturbation can take the form of defective DNA repair or over-compensation of a pathway arising from defects in another pathway^28^. As a result, DNA repair inhibitors could prevent the repair of lesions induced by chemotherapeutic drugs to trigger apoptosis and to enhance the elimination of tumor cells.

We rationalize that patients with high 10-gene scores would have heightened ability for DNA repair thus allowing tumor cells to progress through the cell cycle and continue to proliferate. Using Spearman’s rank-order correlation, we observed that the expression of each of the 10 signature genes were positively correlated with the expression of genes involved in cell cycle progression (cyclins and CDKs) and negatively correlated with genes involved in cell cycle arrest (CDK inhibitors) (Fig. 6A). Interestingly, the patterns of correlation were remarkably similar across all six cancer types, implying that elevated expression of DNA repair genes is associated with a hyper-proliferative phenotype. We next asked whether patients within the high 10-gene score category had an overrepresentation of processes associated with cell cycle dysregulation as this could provide an explanation on the elevated mortality risks in these patients. To answer this, we divided patients from each of the six cancer types into two groups (high score and low score) based on the mean expression of the 10 signature genes using the 50^th^ percentile cutoff. Differential expression analyses between the high- and low-score groups revealed that 394, 425, 1259, 1279, 714 and 977 genes were differentially expressed (−1 > log_2_ fold-change > 1, P<0.05) in clear cell renal cell, papillary renal cell, liver, lung, pancreas and endometrial cancers respectively (Table S4).

**Figure 6.**
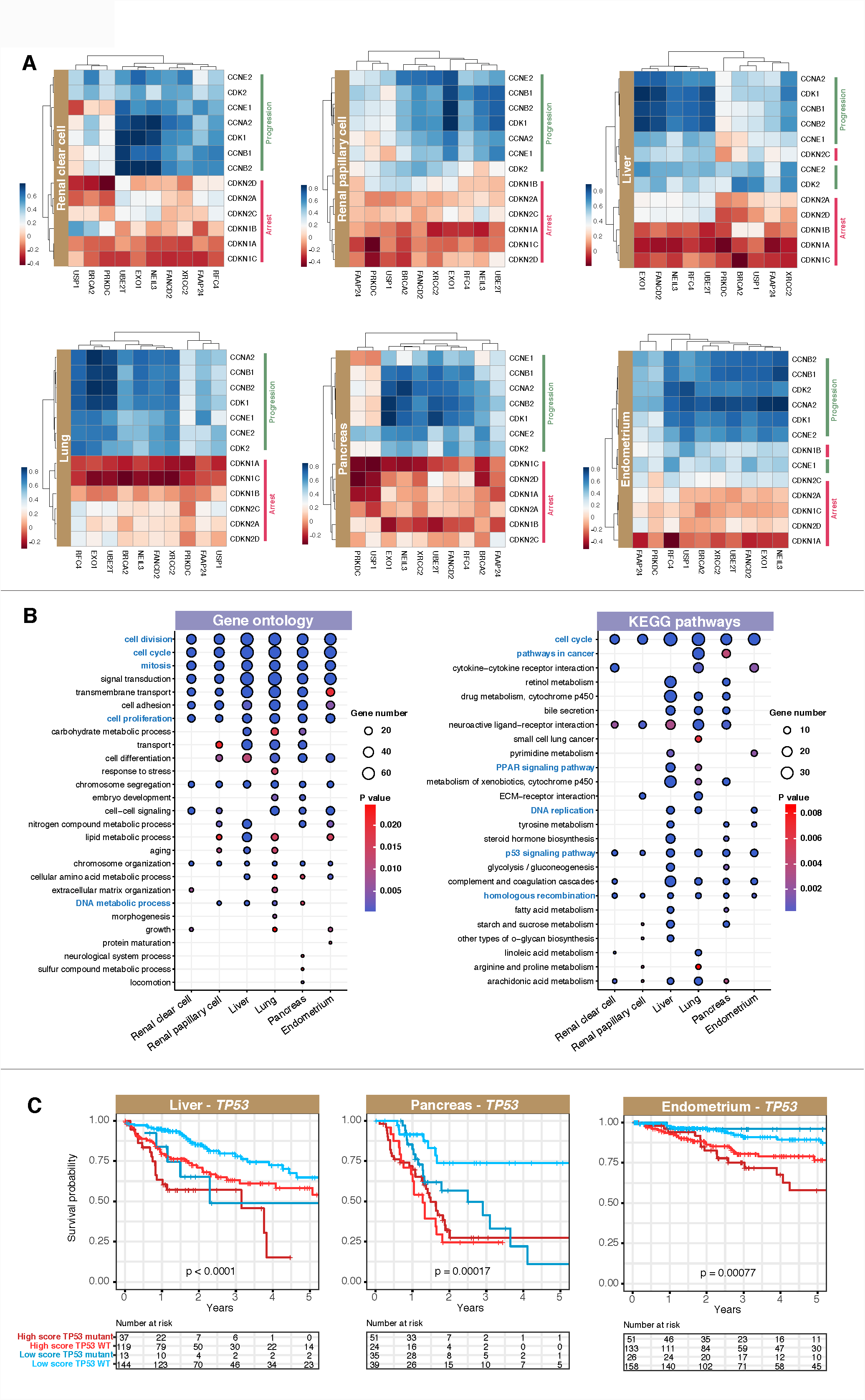
Elevated DNA repair gene expression is associated with an overproliferation phenotype. **(A)** Significant positive correlations between individual signature gene expression and genes involved in cell cycle progression, while negative correlations were observed with genes involved in cell cycle arrest. Heatmaps were generated using the R pheatmap package. Cell cycle genes were depicted on the y-axis and the 10 signature genes on the x-axis. **(B)** Patients were median-stratified into low- and high-score groups using the 10-gene signature for differential expression analyses. Enrichment of GO and KEGG pathways associated with differentially expressed genes were depicted for all six cancers. **(C)** Investigation of the relationship between a gene involved in cell cycle checkpoint regulation, *TP53,* and the signature. Patients were categorized into four groups based on their *TP53* mutation status and 10-gene scores for Kaplan-Meier analyses. P values were determined from the log-rank test.

Analyses of biological functions of these genes revealed functional enrichment of ontologies associated with cell division, mitosis, cell cycle, cell proliferation, DNA replication and homologous recombination consistent in all six cancer types (Fig. 6B). This suggests that the significantly higher mortality rates in patients with high 10-gene scores were due to enhanced tumor cell proliferation exacerbated by the ability of these cells to repair DNA lesions as they arise. Additional ontologies related to tumorigenesis such as *PPAR* and *TP53* signaling were also associated with poor prognosis (Fig. 6B). A total of 87 differentially expressed genes (DEGs) were found to be in common in all six cancer types (Fig. S3) (Table S5). To dissect the underlying biological roles of the 87 DEGs at the protein level, we evaluated the enrichment of transcription factor protein-protein interactions using the Enrichr platform^14^. *TP53* represents the most enriched transcription factor involved in the regulation of the DEGs as evidenced by the highest combined score, which takes into account both Z score and P value (Table S6). This indirectly corroborated our results on enriched *TP53* signaling obtained from the KEGG pathway analysis (Fig. 6B). Taken together, these results highlight the interplay between DDR signaling, cell cycle regulation and *TP53* function in determining prognosis.

### Prognostic relevance of a combined model involving the 10-gene signature and *TP53* mutation status

An important role of *TP53* is its tumor suppressive function through TP53-mediated cell cycle arrest and apoptosis^29^. Hence, somatic mutations in *TP53* can confer tumor cells with growth advantage and indeed, this is a well-known phenomenon in many cancers^30-32^. We rationalized that *TP53* deficiency resulting in defective checkpoint may synergize with the overexpression of DNA repair genes to prevent growth arrest and promote tumor proliferation. To test this hypothesis, we examined *TP53* mutation status in all six cancer types and observed that *TP53* mutation frequency was the highest in pancreatic cancer patients (58%) followed by lung cancer (57%), endometrial cancer (21%), liver cancer (16%), papillary renal cell (1.8%) and clear cell renal cell (1.2%) (Table S7). Cancers with *TP53* mutation frequency of at least 10% were selected for survival analyses. Univariate Cox regression analyses revealed that *TP53* mutation status only conferred prognostic information in pancreatic (HR=1.657, P=0.044), endometrial (HR=1.780, P=0.041) and liver (HR=2.603, P<0.0001) cancers but not in lung cancer (HR=1.428, P=0.056) (Table 1). Cancers where *TP53* mutation offered predictive value were taken forward for analyses in relation to the 10-gene signature. Cox regression analyses revealed that a combination of *TP53* mutation and high 10-gene score resulted in significantly higher risk of death (Table 3; Fig. 6C). Survival rates were significantly diminished in patients harboring high 10-gene scores and the mutant variant of *TP53* compared to those with low 10-gene scores and wild-type *TP53:* liver (HR=3.876, P<0.0001), pancreas (HR=4.881, P=0.0002) and endometrium (HR=3.719, P=0.00028) (Table 3; Fig. 6C). Moreover, in multivariate Cox models involving TNM staging and *TP53* mutation status, the 10-gene signature remained a significant prognostic factor (Table 1). This suggests that although the 10-gene signature provided additional resolution in risk assessment when used in combination with *TP53* mutation status, its function is independent. However, in the multivariate model *TP53* was significant only in liver cancer (HR=2.085, P=0.0044), suggesting that *TP53* mutation was not independent of the signature or TNM staging in pancreatic and endometrial cancers (Table 1). Overall, the results suggest that defects in cell cycle checkpoint combined with augmented DNA repair ability were adverse risk factors contributing to poor prognosis. Both *TP53* mutation status and 10-gene scores could offer additional predictive value in risk assessment by further delineation of patients into additional risk groups.

## Discussion and Conclusion

We systematically examined the associations between the expression patterns of 138 DNA repair genes in 16 cancer types and prognosis. Our pan-cancer multigenic approach revealed genes that work synergistically across cancers to inform patient prognosis that would otherwise remain undetected in analysis involving a single gene or a single cancer type. We developed a 10-gene signature that incorporates the expression profiles of 10 highly correlated DNA repair genes for use as risk predictors in six cancer types (n=2,257). This signature offers a more precise discrimination of patient risk groups in these six cancers where high expression of signature genes is associated with poor survival outcomes. Importantly, we demonstrated that the signature can improve the prognostic discrimination of TNM when used as a combined model, which is particularly useful to allow further stratification of patients within similar TNM stage groups (Fig. 4).

Intrinsic differences in DNA repair machineries in cancer cells may pose a significant challenge to successful therapy. Mutations in DNA repair genes allow the generation of persistent DNA lesions that would otherwise be repaired. Germline mutations of DNA repair genes are linked to increased genome instability and cancer risks^33^ and abrogation of genes in one DNA repair pathway can be compensated by another pathway^1^. *BRCA1* and *BRCA2* mutations sensitize cells to PARP1 inhibition, a protein involved in the BER pathway^10^. Since *BRCA1* and *BRCA2* are important for homology-directed repair, PARP1 inhibition in BRCA1/2-defective cells would result in dysfunctional HDR and BER pathways preventing lesion repair and thus leading to apoptosis^10^.

In addition to genetic polymorphism, upregulation of DNA repair genes in tumors could promote resistance to radiotherapy and chemotherapy as the cells would have enhanced ability to repair cytotoxic lesions induced by these therapies. Overexpression of *ERCC1* involved in the NER pathway in non-small-cell lung cancer is linked to poor survival in cisplatin-treated patients^7^. The 1,2-d(GpG) cross-link lesion generated by cisplatin treatment is readily repaired by the NER pathway, hence *ERCC1* overexpression would promote cisplatin resistance. Low *MGMT* expression in astrocytoma is associated with longer survival outcomes in patients treated with temozolomide^34^; an observation that is consistent with the role of *MGMT* in repairing lesions caused by temozolomide thus allowing *MGMT* deficient tumor cells to accumulate enough unrepairable damage. *TP53* plays essential roles in cell-cycle arrest and apoptosis through the activation of checkpoint genes^29^. We show that patients with high 10-gene scores that concurrently have mutant *TP53* exhibited significantly higher mortality rates (Fig. 6C), suggesting that defects in cell cycle checkpoint coupled with an increase propensity for DNA repair may lead to dramatically poorer outcomes.

Multiple studies have reported the associations between dysfunctional DNA repair pathways and cancer, but most of these studies are restricted to investigations on a limited number of genes and on one cancer at a time. One of the key advantages of our study is that it is an unbiased exploration transcending the candidate-gene approach that takes into account the multifaceted interplay of DNA repair genes in diverse cancer types. We rationalize that since ionizing radiation and chemotherapy are the main treatment options currently available for cancer patients, a molecular signature capable of discriminating patients with increased expression of DNA repair genes that would benefit from adjuvant therapy through pharmacological inhibition of DNA repair to overall improve therapeutic outcomes.

Tumor hypoxia is also a well-known cause of therapy resistance. A notable finding of our study is that patients having both high 10-gene and hypoxia scores had significantly poorer survival rates compared to those with low 10-gene and hypoxia scores (Fig. 5). Previous reports suggest that low oxygen conditions may interfere with DNA damage repair. For example, hypoxia could compromise HR function through decreased *RAD51* expression^35^. However, results concerning the effects of hypoxia on DDR signaling have remained inconclusive. Genes associated with NHEJ were reported to be downregulated under hypoxia in prostate cancer cell lines^36^, while hypoxia drove the upregulation of NHEJ-associated genes, *PRKDC*and *XRCC6,* in hepatoma cell lines^37^. The authors proposed an interaction between *PRKDC* and the hypoxia-responsive transcriptional activator, HIF-1α, hence suggesting that tumor hypoxia may lead to increase in NHEJ. Tumor cells within their 3D space are subjected to differential levels of oxygen over time and chronic exposures to these fluctuating conditions could result in very different biological outcomes. *In vitro* studies retain a significant caveat as many hypoxia assays are carried out short term using constant, predefined oxygen tensions. Although further work is needed to ascertain the clinical relevance of these findings, our results clearly demonstrate that the integration of hypoxia assessment in molecular stratification using the 10-gene signature revealed a subset of high-risk individuals accounting for approximately 31% to 38% in each cohort (Fig. 5B). Whether hypoxia could directly promote DNA damage repair *in vivo* remains an open question.

We reasoned that the expression patterns of DNA repair genes would positively correlate with genes involved in cell cycle progression since lesions could be repaired more effectively to prevent cell cycle arrest (Fig. 6A). Enhanced DNA repair ability may also confer tumor cells with growth advantage. Consistent with this hypothesis, differential expression analyses between patients with high versus low 10-gene scores revealed an enrichment of ontologies involved in growth stimulation as a consequence of increased DNA repair gene expression (Fig. 6B). Enrichment of biological pathways involved in cell cycle, mitosis, cell division and DNA replication implied that the shorter life expectancy in patients with high 10-gene scores could in part be explained by an overproliferation phenotype commonly present in more aggressive tumors.

In summary, we developed a prognostic signature involving DNA repair genes and confirmed its utility as a powerful predictive marker for six cancer types. Although not currently afforded by this work due to its retrospective nature, it will be useful to determine if the signature can predict response to radiotherapy and chemotherapy in future research. While prospective validation is warranted, we would expect, based on our encouraging retrospective data, that the signature can guide decision making and treatment pathways. The confirmation of this hypothesis by a clinical trial using the 10-gene signature to select patients that would benefit from treatment with adjuvant DNA repair inhibitors could have a substantial impact on treatment outcomes.

## Conflict of Interest

None declared.

## Funding

None.

## Authors contribution

WHC and AGL designed the study, analyzed the data and interpreted the data. AGL supervised the research. WHC and AGL wrote the initial manuscript draft. AGL revised the manuscript draft and approved the final version.

## Supplementary information

**Figure S1.**
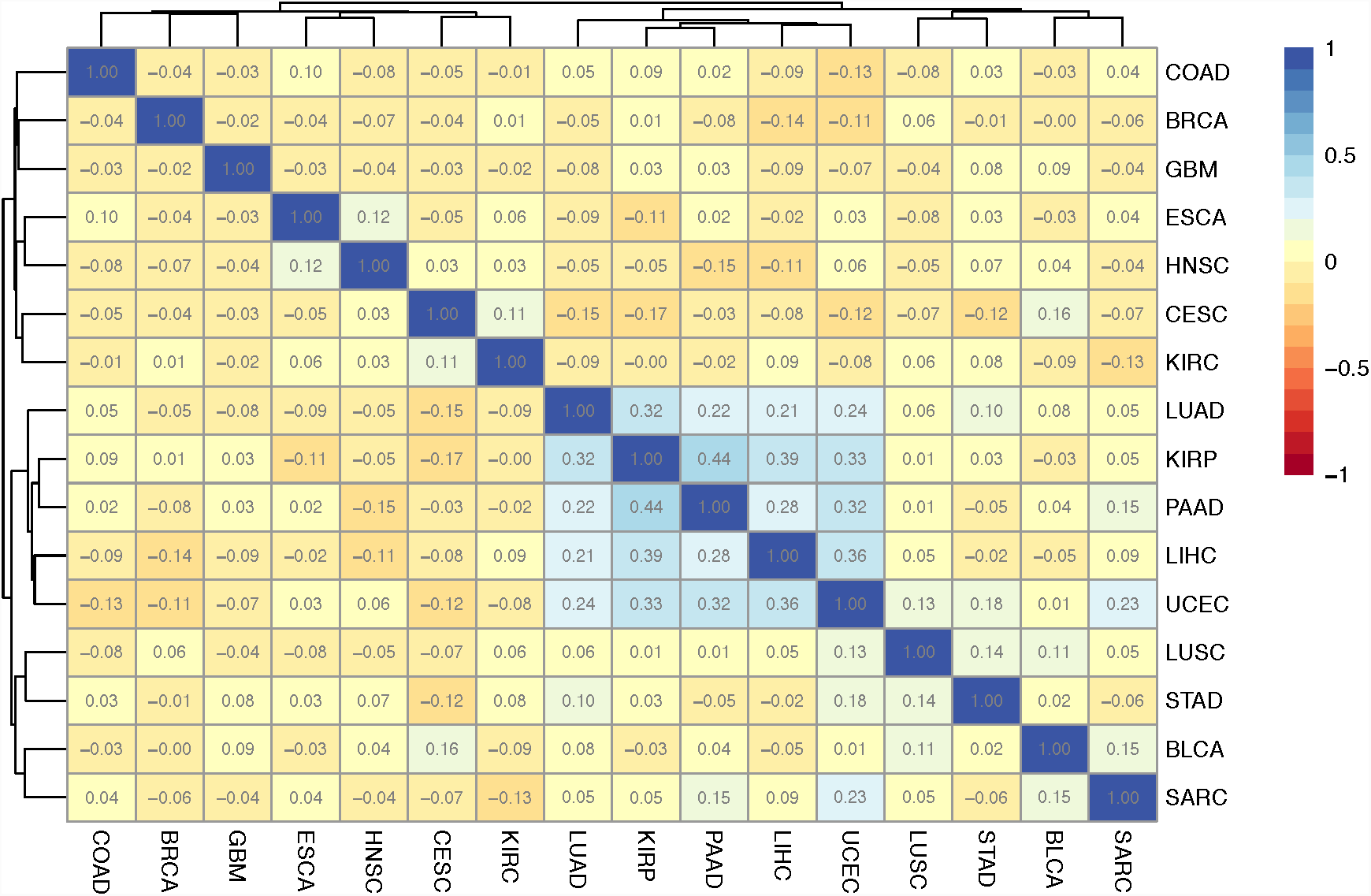
Correlation analyses of 138 prognostic DNA repair genes. Spearman’s correlation coefficients were determined from pairwise comparisons prognostic genes from 16 cancer types. Five cancers were highly correlated as shown in the blue area of the heatmap. Numbers represent correlation coefficient values. Refer to Table S2 for cancer abbreviations.

**Figure S2.**
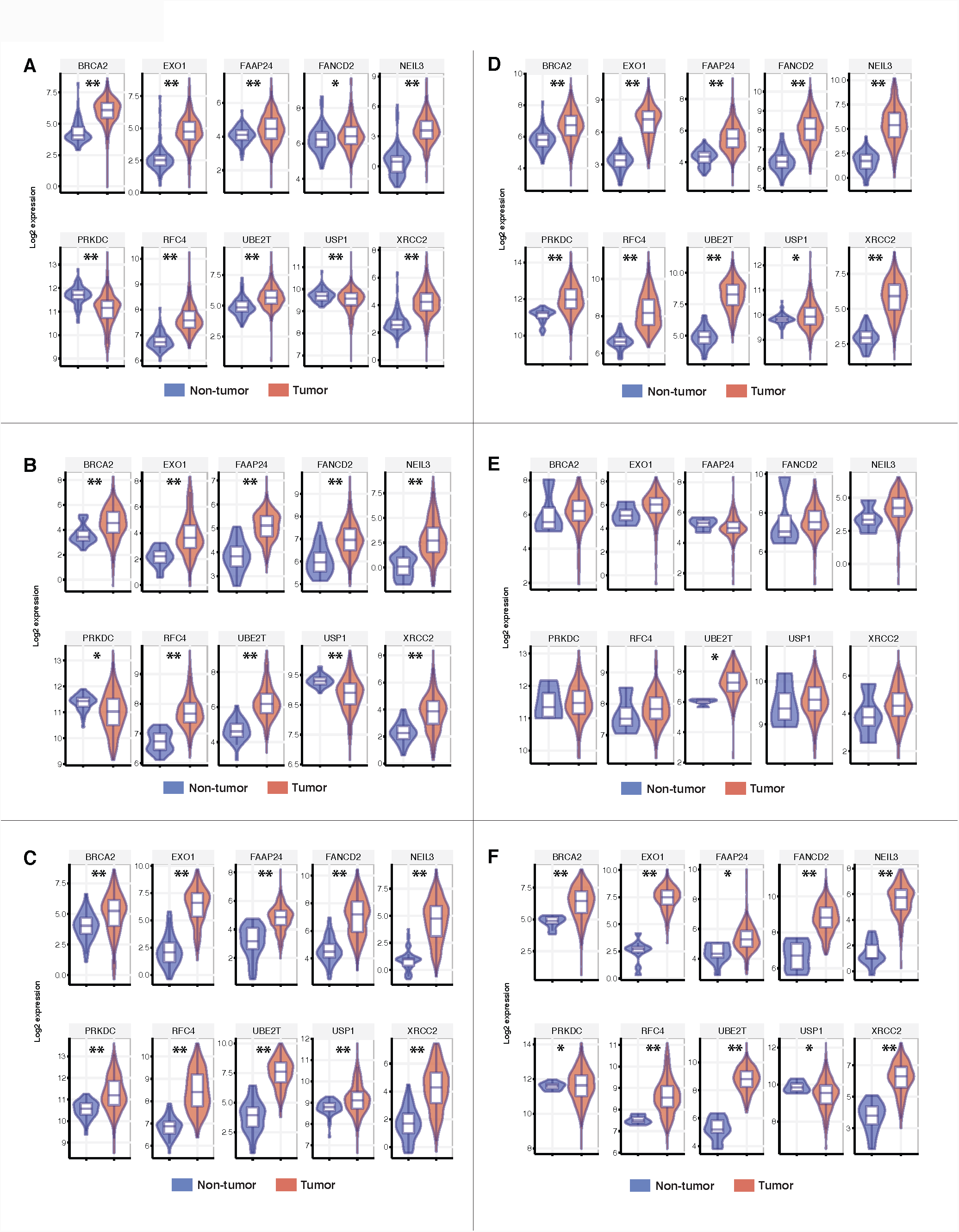
Expression distribution of the ten signature genes in tumor and non-tumor samples. Boxplots overlaying violin plots were used to illustrate tumor and non-tumor distribution in six cancers: **(A)** clear cell renal cell, **(B)** papillary renal cell, **(C)** liver, **(D)** lung, **(E)** pancreas and **(F)** endometrium. Nonparametric Mann-Whitney-Wilcoxon tests were employed to determine whether there were significant differences in expression distributions. Asterisks represent significant P values: * < 0.05, *** < 0.0001.

**Figure S3.**
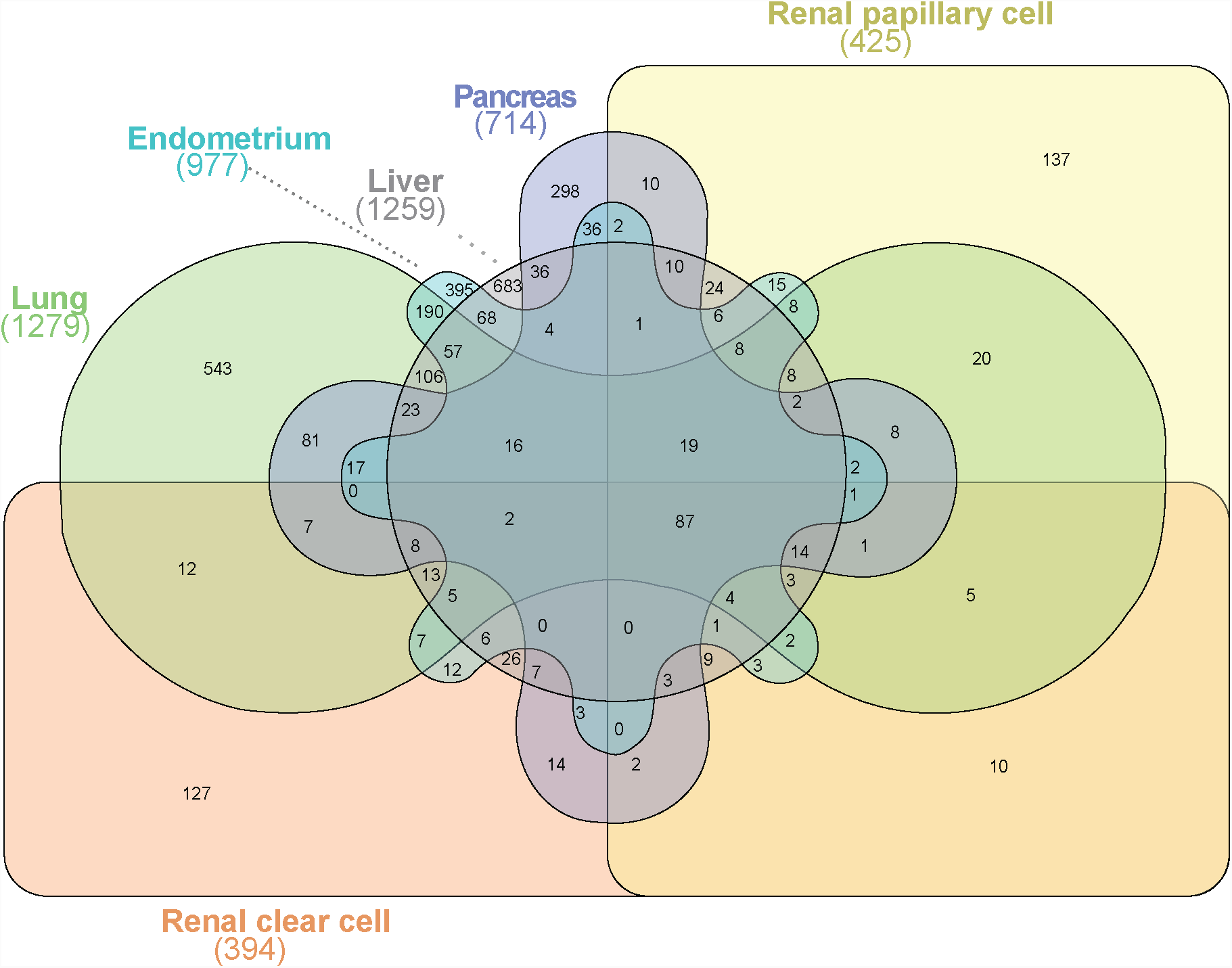
Venn diagram depicts a six-way comparison of the differentially expressed genes (-1 > log_2_ fold-change > 1, P<0.05) identified from high-score versus low-score patients in all six cancers. Numbers in parentheses represent the number of differentially expressed genes in each cancer. The Venn intersection of all cancers indicated that 87 genes were common.

**Table 1.** Univariate and multivariate Cox proportional hazards analyses of the 10-gene signature and additional clinical risk factors associated with overall survival in six cancers.

**Table 2.** Univariate Cox proportional hazards analysis of the relation between the 10-gene signature and hypoxia score.

**Table 3.** Univariate Cox proportional hazards analysis of the relation between the 10-gene signature and *TP53* mutation status.

**Table S1.** List of 138 DNA repair genes and associated pathways.

**Table S2.** Description of TCGA cancer cohorts.

**Table S3.** Univariate Cox proportional hazards analysis of the 138 genes in 16 cancers.

**Table S4.** Differentially expressed genes between high- and low-score patient groups in six cancers.

**Table S5.** List of 87 differentially expressed genes that are common in all six cancers.

**Table S6.** Enrichr transcription factor protein-protein interaction analysis of the 87 differentially expressed genes.

**Table S7.** *TP53* mutation analysis in liver, pancreatic, endometrial and lung cancers.

## List of abbreviations

DDR: DNA damage response
BER: Base excision repair
NER: Nucleotide excision repair
MR: Mismatch repair
HDR: Homology-directed repair
NHEJ: Non-homologous end joining
FA: Fanconi anemia
TCGA: The Cancer Genome Atlas
GO: Gene Ontology
KEGG: Kyoto Encyclopedia of Genes and Genomes
HR: Hazard ratio
ROC: Receiver operating characteristic
AUC: Area under the curve
TNM: Tumor, node and metastasis
CDK: Cyclin-dependent kinase
DEG: Differentially expressed genes

